# MASH: Mediation Analysis of Survival Outcome and High-dimensional Omics Mediators with Application to Complex Diseases

**DOI:** 10.1101/2023.08.22.554286

**Authors:** Sunyi Chi, Christopher R Flowers, Ziyi Li, Xuelin Huang, Peng Wei

## Abstract

Environmental exposures such as cigarette smoking influence health outcomes through intermediate molecular phenotypes, such as the methylome, transcriptome, and metabolome. Mediation analysis is a useful tool for investigating the role of potentially high-dimensional intermediate phenotypes in the relationship between environmental exposures and health outcomes. However, little work has been done on mediation analysis when the mediators are high-dimensional and the outcome is a survival endpoint, and none of it has provided a robust measure of total mediation effect. To this end, we propose an estimation procedure for <u>M</u>ediation <u>A</u>nalysis of <u>S</u>urvival outcome and <u>H</u>igh-dimensional omics mediators (MASH) based on sure independence screening for putative mediator variable selection and a second-moment-based measure of total mediation effect for survival data analogous to the *R*^2^ measure in a linear model. Extensive simulations showed good performance of MASH in estimating the total mediation effect and identifying true mediators. By applying MASH to the metabolomics data of 1919 subjects in the Framingham Heart Study, we identified five metabolites as mediators of the effect of cigarette smoking on coronary heart disease risk (total mediation effect, 51.1%) and two metabolites as mediators between smoking and risk of cancer (total mediation effect, 50.7%). Application of MASH to a diffuse large B-cell lymphoma genomics data set identified copy-number variations for eight genes as mediators between the baseline International Prognostic Index score and overall survival.

## 1. Introduction

Mediation analysis is performed to elucidate how an exposure influences an outcome through mediating variables. Researchers have used this approach widely in epidemiology and clinical studies, for example, to investigate the connection between environmental risk factors and disease outcomes via intermediate molecular phenotypes, such as gene expression or protein markers. To date, most mediation studies have focused on a single mediator or a few mediators, with much less attention paid to high-dimensional mediators [Daniel et al. (2015); VanderWeele et al. (2014)], particularly in the analysis of survival data [Huang et al. (2017)]. Mediation analysis of high-dimensional data has garnered increasing interest as a result of rapid technological advancements in high-throughput genomic profiling such as RNA sequencing. Although researchers have recently made progress in high-dimensional mediation analysis of continuous outcomes in the linear model setting [Liu et al. (2021); Sampson et al. (2018)], investigators have devised only a few methods for high-dimensional mediation analysis of survival data owing to the considerable challenges in variable selection, censoring, and estimation of the total mediation effect. Luo et al. (2020) proposed a procedure called HIMA for identifying the putative mediators and estimating the indirect effects of DNA methylation on the pathway from smoking to overall survival among lung cancer patients in the Cancer Genome Atlas (TCGA) lung cancer cohort. HIMA used the classic product-based mediation effect measure that could cancel the component-wise mediation effects in a multiple-mediator model owing to mediation direction disagreement, which is commonly encountered in high-dimensional genomic studies. In fact, real data application results of HIMA showed that the indirect effect by the product measure was negative due to the cancellation of both positive and negative component-wise mediation effects, resulting in that the total effect was less than the direct effect. In addition, the classic product-based and difference-based total mediation effect measures for linear models are only applicable in survival models for rare outcomes under the counterfactual framework [VanderWeele et al. (2011)].

As an alternative to the first-moment-based product effect measure, Fairchild et al. (2009) first proposed to use the *R*^2^-based mediation effect size measure for a single mediator in the linear model setting. Recently, Yang et al. (2021) extended the use of *R*^2^ measure to multiple- or high-dimensional-mediator models for a continuous outcome. Importantly, Yang et al. (2021) showed that the *R*^2^ measure can capture the nonzero total mediation effect in the presence of bi-directional component-wise mediation effects, which are likely ubiquitous in high-dimensional omics data settings. Shi et al. (2022) further extended the *R*^2^ measure to survival outcomes and compared five *R*^2^ measures in a Cox proportional hazards model owing to the non-unique definition of *R*^2^ in survival models. However, to the best of our knowledge, the *R*^2^ measure has yet to be studied with high-dimensional mediator models for survival outcomes, which are common in prospective cohort studies, such as the Framingham Heart Study (FHS), and clinical studies, such as the TCGA project.

To fill this gap in knowledge described above, we propose the MASH method: <u>M</u>ediation <u>A</u>nalysis of <u>S</u>urvival outcome and <u>H</u>igh-dimensional omics mediators based on a three-step mediator selection procedure and the second-moment mediation effect size measure for survival models analogous to the *R*^2^ measure in a linear model. In addition to extending the *R*^2^ measure to high-dimensional survival data, we propose the use of a partial *R*^2^ mediation measure to adjust for confounding effects.

To evaluate our mediation effect estimation in terms of bias and variance, as well as mediator selection accuracy, we conducted extensive simulation studies in various settings by varying the censoring rate, the number of putative mediators and the directions of mediation effects. We demonstrated that our approach has better performance than existing methods in terms of bias and variance in mediation effect estimation and is stable to data with high censoring rate. Furthermore, we applied the MASH to the metabolomics data of 1919 subjects in the FHS and a diffuse large B-cell lymphoma (DLBCL) genomics data set to address three problems: how smoking influences the risk of coronary heart disease through metabolites, how smoking influences the risk of cancer through metabolites, and how the baseline International Prognostic Index (IPI) impacts overall survival of DLBCL through copy-number variations.

## 2. Methods

Mediation processes are framed in terms of intermediate variables between an independent variable and a dependent variable, with a minimum of three variables required in total. Let *T* denote the dependent variable – a survival outcome, *X* denote the independent variable, an exposure of interest and *M* denote the set of mediator variables that is supposed to transmit the effect of *X* to *T* . Figures 1A and 1B illustrate the relationships among the survival outcome, exposure, and mediators.

**FIG 1.**
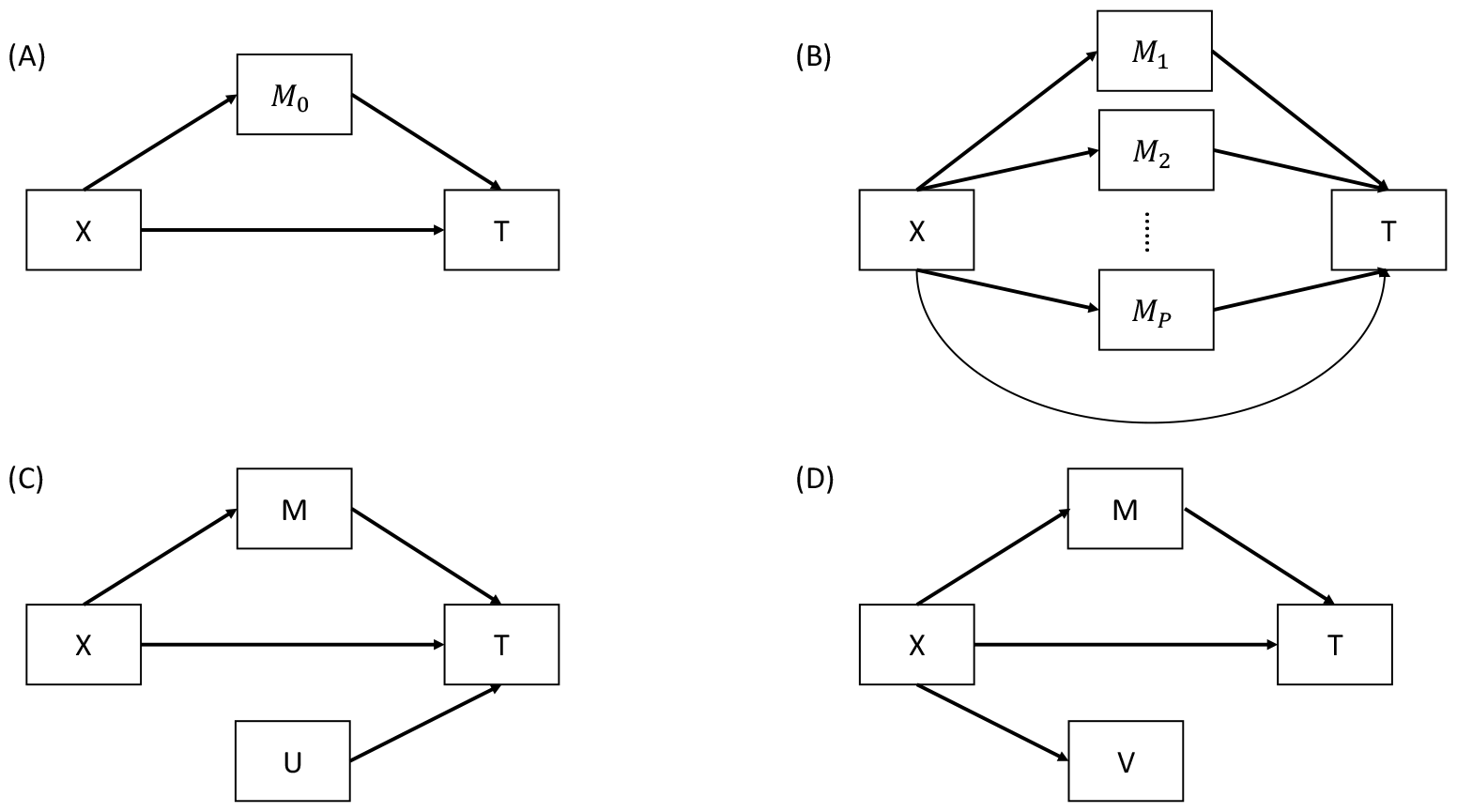
Diagrams for potential mediation settings. *X* is the independent variable, *T* is the survival outcome, and *M* is a set of true mediator. *U* is a set of variables associated with *Y* but not *X*, and *V* is a set of variables associated with *X* but not *Y* . (A) A single-mediator model. (B) A multiple-mediator model. (C) A model with non-mediator *U* . (D) A model with non-mediator *V* .

In our context in which all the mediators are measured at the same time and a direct causal relationship among them is less likely to exist, we assume that the mediators are not causally related to each other. Although we will propose a method to control for confounding variables in section 2.3, the following assumptions are made as in VanderWeele et al. (2014): (1) no unmeasured confounders between the exposure and the survival outcome, between the mediators and the survival outcome or between the exposure and the mediators, and (2) no exposure-induced confounding between the mediators and the survival outcome.

### 2.1. Review of Mediation Models

We use the following Cox proportional hazards regression models to assess the role of mediators M in the pathway from exposure X to survival outcome T:

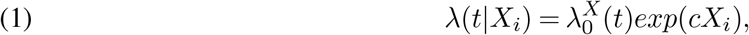

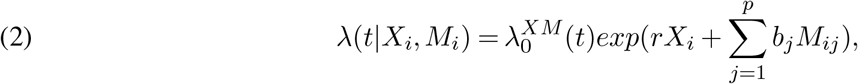

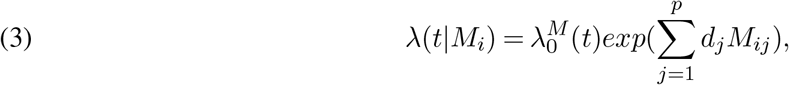

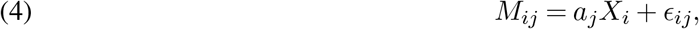

where *M*_*i*_ = (*M*_*i*1_, *M*_*i*2_, … , *M*_*ip*_)^*′*^ is the *p*-dimensional mediator vector for subject *i* = 1, 2, 3, …, *n* and, without loss of generality, *X* and *M*_*j*_ are standardized to have mean 0 and variance 1. Let *D* denote the time to event and *C* denote the censoring time. The observed survival outcome is *T*_*i*_ = *min*(*D*_*i*_, *C*_*i*_), and the failure indicator is *δ*_*i*_ = *I*(*D*_*i*_ *≤ C*_*i*_) for *i* = 1, 2, 3, …, *n*. Equations (1), (2), and (3) are Cox proportional hazards models describing the relationship (1) between *T* and *X*, (2) between *T* , *X*, and *M* , and (3) between *T* and *M* , respectively.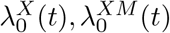, and 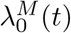 are their respective baseline hazard functions. In a Cox proportional hazards regression model, the hazard ratio is usually used as a measure of an independent variable’s effect on a dependent variable. In equations (1), (2), and (3), *c* is the parameter relating the exposure to the outcome; *r* is the direct effect parameter relating the exposure to the outcome; *b*, defined as (*b*_1_, … , *b*_*p*_)^*T*^ , is the parameter vector relating the mediators to the outcome with adjustment for the effect of the exposure; and *d* = (*d*_1_, … , *d*_*p*_)^*′*^, is the parameter vector relating the mediators to the outcome. Equation (4) characterizes how exposure influences the mediators, where *a* = (*a*_1_, … , *a*_*p*_)^*′*^ is the parameter vector relating the exposure to the mediators and residual *ϵ*_*i*_ = (*ϵ*_*i*1_, *ϵ*_*i*2_, … , *ϵ*_*ip*_) *∼ MV N* (0, *E*_*p×p*_), for *i* = 1, … , *n*.

To formally define the causal effect for Cox model in counterfactual framework [Robins and Greenland (1992); Pearl (2001)], we follow the framework of VanderWeele et al. (2011) and Huang et al. (2017) using difference in log-hazard scale. Let *λ*^*x*,*M*(*x*)^(*t*) denote the hazard function of *T*_*x*,*M* (*x*)_ when *X* was set to *x* and *M* was set to the value it would have taken if *X* was set to *x*. Let *λ*^*x∗*,*M* (*x∗*)^(*t*) denote the hazard function of *T*_*x∗*,*M* (*x∗*)_when *X* was set to *x*^*∗*^ and *M* was set to the value it would have taken if *X* was set to *x*^*∗*^. Thus, the natural direct effect (NDE) is defined as the difference between *logλ*^*x∗*,*M* (*x*)^(*t*) and *logλ*^*x*,*M*(*x*)^(*t*), and the natural indirect effect (NIE) is defined as the difference between *logλ*^*x∗*,*M* (*x∗*)^(*t*) and *logλ*^*x∗*,*M* (*x*)^(*t*). The total effect (TE) of *X* on *T* can then be decomposed into two components: a direct effect of *X* on *T* (NDE) and an indirect effect of *X* on *T* through *M* (NIE). Under the rare outcome assumption, the NDE approximate to (*x*^*∗*^ *− x*)*r*, the NIE approximate to 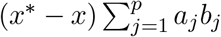 , and the TE approximate to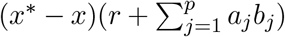. Without loss of generality, we set *x*^*∗*^ = 1 and *x* = 0. Therefore, 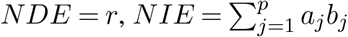, and 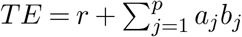.

### 2.2. Measures of mediation effects

The standard difference measure (i.e., *c − r*) and product measure 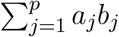for the indirect effect coincide for a continuous outcome with *c, r, a*_*j*_, *b*_*j*_ similarly defined in a linear model as in models (1), (2) and (3) (MacKinnon (2008)). They are also comparable based on simulations in single-mediator analysis without considering censoring and ties in log survival time models and log hazard time models, but not in Cox proportional hazards models [Tein and MacKinnon (2003)]. In single-mediator Cox model, the difference method uses *c − r* as a measure of the indirect effect and the product method uses *ab*. VanderWeele et al. (2011) showed that if all of the models are correctly specified and the outcome is rare, these two are approximately equal in single-mediator Cox model. However, neither the product measure nor the difference measure for the proportional hazards model has any sort of clear causal interpretation as a measure of mediation effect VanderWeele et al. (2011). In multiple-mediator Cox models, the difference and product measures are, respectively, defined as *c − r* and Σ *a*_*j*_*b*_*j*_, which are approximately equal only under rare events/high censoring rates (VanderWeele et al. (2011),Shi et al. (2022)). Furthermore, in high-dimensional mediation analysis the component-wise mediation effects (*a*_*j*_*b*_*j*_’s) could cancel out due to the likely presence of opposite mediation directions, resulting in close to zero or negative product measure. However, existing mediation analysis methods for multiple mediators and survival outcomes still use the product measure. Luo et al. (2020) proposed a high-dimensional mediation analysis framework for survival outcomes using the product measure that led to negative product measure and the the estimated total effect less than the direct effect in their real data applications. On the other hand, the *R*^2^ mediation effect measure, which was originally proposed in a single-mediator model by Fairchild et al. (2009), does not suffer from the issue of misleading cancellation when the mediation effects have both positive and negative components. Also, it has good performance with complex structured data, such as highly correlated mediators (Yang et al. (2021)). Therefore, here we propose a explained-variation measure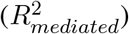 based on second moments to estimate mediation effects of high-dimensional mediators for survival outcomes. This measure does not require rare event assumption and has more robust estimation in multi-mediator analysis than the standard product measure as to be shown later on.

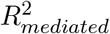 , the measure of mediation effect, is defined by the amount of variation in outcome *T* that is explained by exposure *X* through mediators *M* . In Cox proportional hazards models, this measure is computed as follows: (5)

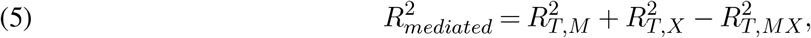

Where 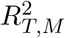 is the proportion of variation in outcome *T* that is explained by the mediators, 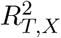 is the variation in *T* that is explained by the exposure, and 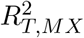 is the proportion of variation in *T* that is explained by both the mediators and the exposure. Thus, 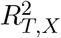 quantifies the total effect whereas 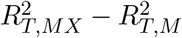 is the direct effect. And the *R*^2^ measure of indirect effect is the total effect minus the direct effect, which is given in equation (5). The shared over simple effect (SOS), defined as 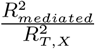 by Lindenberger and Pötter (1998) is the standardized exposure-related variation in the outcome that is shared with the mediators. Although *R*^2^ is clearly defined in linear regression models, applying the notion of explained variation to survival data is not straightforward because of nonnormality of the errors and censoring of the dependent variable. There have been many versions of *R*^2^-like measures proposed for survival data. Royston (2006) outlined the properties that a good explained variation measure for survival models should have: 1) approximate independence of the amount of censoring; 2) reduction to (or a close relationship with) the *R*^2^ usually obtained via “equivalent” linear regression analysis of the same data set, if possible; 3) the nesting property for two models: M1 *⊂* M2 (with *⊂* denoting nesting) and *R*^2^(*M* 1) *≤ R*^2^(*M* 2); 4) increase in *R*^2^ as the strength of the association increases; and 5) availability of confidence intervals. Shi et al. (2022) compared five measures of explained variation for survival data in mediation models and suggested the use of 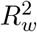 by Kent and O’Quigley (1988) in multiple-mediator analysis in the Cox proportional hazards models. Specifically, 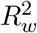 is defined as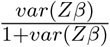, where *Z* is the vector of independent variables and *β* is the coefficient vector in a Cox model. Plugging 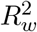 in equation (5) results in the 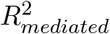 for our proposed high-dimensional mediation analysis for survival outcomes.

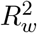 is an approximation of explained variation used to measure the information gain [Kent and O’Quigley (1988)]. It is compatible with the *R*^2^ measure in linear regression models and satisfies the properties out-lined by Shi et al. (2022). Moreover, Schemper and Stare (1996) noted that 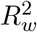 is unaffected by censoring and has good performance across simulation settings comparing measures of explained variation in survival analysis. Importantly, 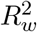 is robust to use in our proposed high-dimensional mediation analysis framework based on our simulation studies and real data applications.

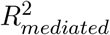 can be shown as a function of *a, b, c, d*, and *r* as follows:

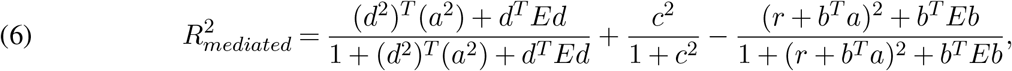

where 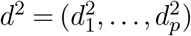 and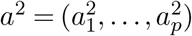.

In the scenario of all *a*’s and *b*’s equal to 0, both product measure and 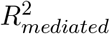 are equal to 0. Of note, 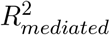 could be non-zero even if 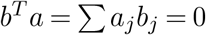 but not all *a*’s and *b*’s are 0. The derivation of equation (6) and comparison of 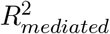 and the product measure in no indirect effect scenarios is presented in the supplementary materials Section 2.

### 2.3. New method

*MASH* To perform mediation analysis of high-dimensional omics mediators such as metabolomics and genome-wide copy-number alterations, we must select the true mediators in the pathway from the exposure to the outcome (Fig. 1B). Let **S**_**0**_ denote a set of potential mediators, **M** denote a set of true mediators, **U** denote a set of variables associated with the outcome but not the exposure (Fig. 1C), and **V** denote a set of variables associated with the exposure but not the outcome (Fig. 1D). None of the **U** or **V** variables are true mediators. Although inclusion of U in mediation analysis for continuous outcomes does not affect the estimation of the *R*^2^ mediation effect [Yang et al. (2021)], our proof in Supplementary Material Section 2 and results of our preliminary simulation study shown in Supplementary Material Table S4 demonstrated that inclusion of either *U* or *V* could bias the estimation of the *R*^2^ mediation effect on survival outcomes. Therefore, we propose a three-step mediator selection procedure as illustrated in Figure 2 to identify true mediators between the exposure and outcome by excluding **U, V**, and noise variables (i.e., those not associated with the exposure or the outcome).

**FIG 2.**
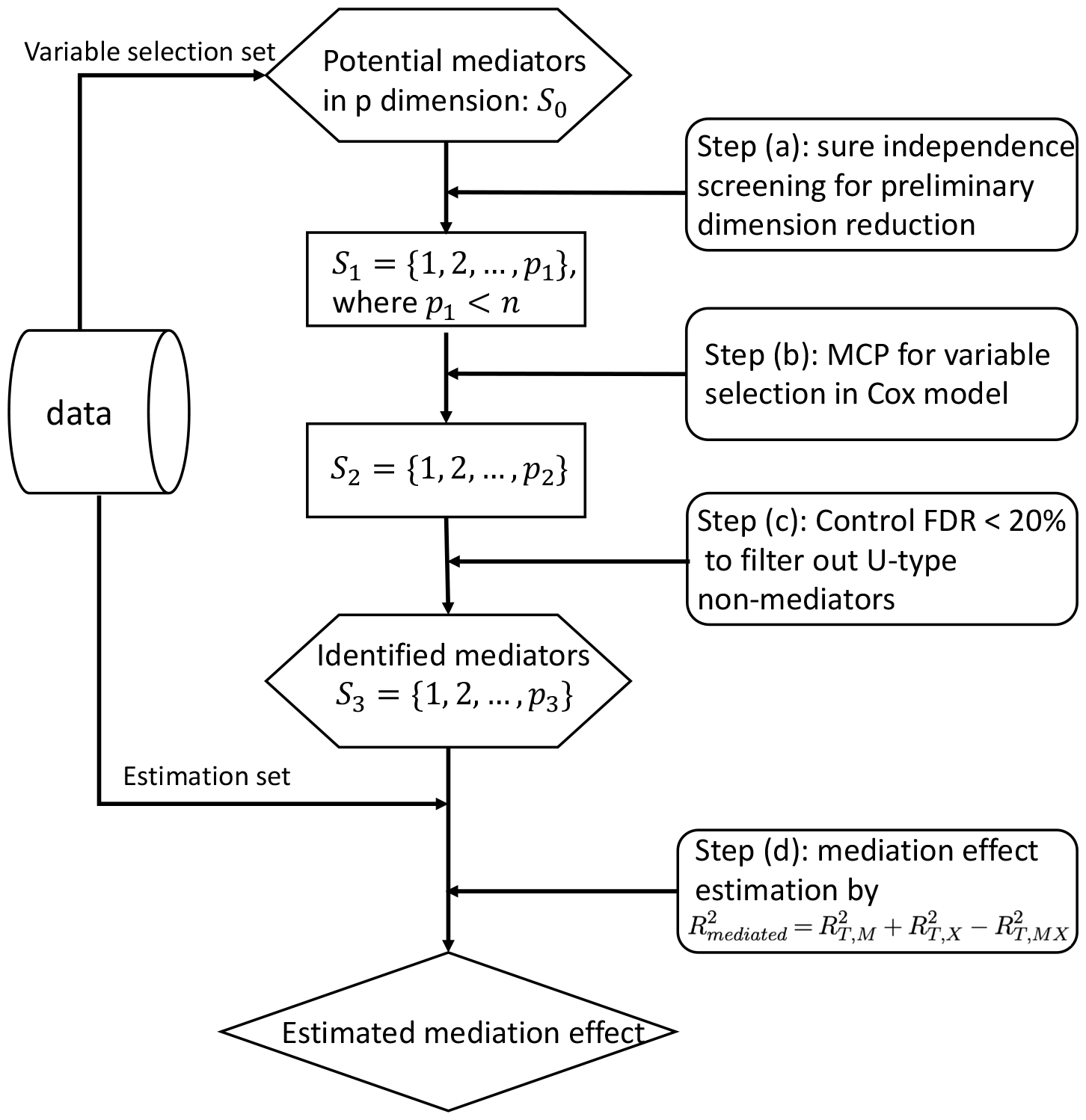
Overall workflow of MASH. The workflow consists of four main steps: (a) sure independence screening for preliminary screening, (b) MCP for variable selection, (c) FDR control at 0.2 level for variable selection, and (d) mediation effect estimation using *R*^2^-like measurements.

Traditional statistical methods fail when the number of mediators *p* is much larger than the sample size *n*. Sure independence screening is a large-scale screening method for an ultra-high dimensional feature space with the property that all the true variables survive after variable screening with the probability tending to 1 [Fan and Lv (2008)]. Principled sure independence screening by Zhao and Li (2012) is an extension of sure independence screening for censored survival data using the Cox proportional hazards model, which resembles the marginal ranking method proposed by Fan et al. (2010). This method avoids the requirement of choosing a size of subset to retain after screening by specifying the desired false-positive rate. Therefore, as shown in Figure 2, with MASH, principled sure independence screening based on Cox proportional hazards model (3) is first used to reduce the dimension and exclude *V* -type non-mediators and noise variables. In this step, a subset of potential mediators *S*_1_ is selected under a fixed false-positive rate of 0.05; those are the mediators likely to be associated with the outcome. Variable selection within the subset *S*_1_ via the minimax concave penalty (MCP) is then conducted to further exclude putative mediators not associated with the outcome based on the Cox proportional hazards model (3). Ten-fold cross validation for MCP-based survival models is used over a grid of values for the regularization parameter *λ*. Next, the false discovery rate (FDR) adjusted *p* value based on the Benjamini-Hochberg procedure with a critical value of 0.2 for the model in equation (4) is used to test the marginal association of each selected potential mediator with the exposure to exclude *U* -type non-mediators and noise variables. Finally, the mediation effect 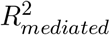 and SOS are estimated in the estimation dataset using the selected mediators in the variable selection dataset. Of note, as shown in Figure 2, the data are split into two halves with one half used as a variable selection set to select the true mediators and the other half used as an independent dataset to estimate the mediation.

### 2.4. Partial R^2^ to control confounding effects

As illustrated by VanderWeele (2016) failure to control for exposure-outcome, exposure-mediator and mediator-outcome confounding effects in mediation analysis can substantially bias estimates of mediation effects, whereas adjusting for measured confounders as covariates in mediation models can reduce the bias. Therefore, we propose a partial *R*^2^ measure to adjust for potential confounders in mediation analysis. Let *Z* denote baseline covariates such as age, which may act as a potential confounder. Our models adjusting for confounders are as follows:

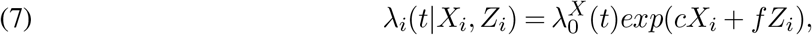

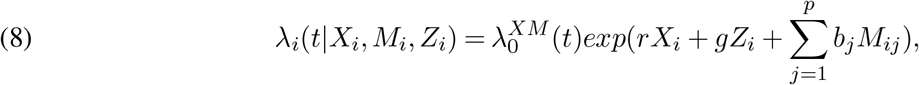

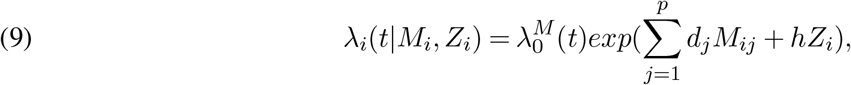

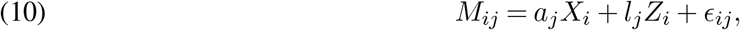

In addition to controlling for mediator-outcome confounding effects, we also propose models adjusting for exposure-outcome confounders, which include equations (4), (7), (8), (9), and (11) as follows:

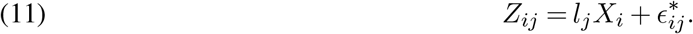

In general, partial *R*^2^ is defined as the proportion of variation in the outcome variable that cannot be explained in a reduced model but can be explained by the predictors specified in a full model. To adjust for the covariates in the indirect effect of exposure on the outcome through mediators, partial *R*^2^ is adapted based on our original formulas to calculate the mediation effect as shown below.

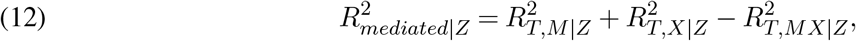

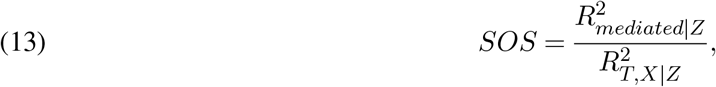

where, according to the definition of partial 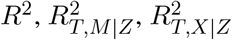 , and 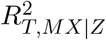 are

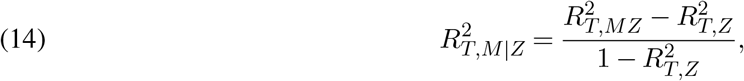

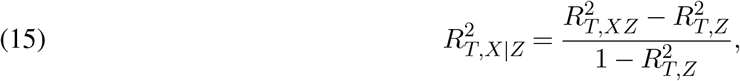

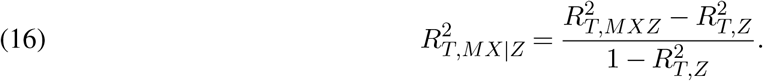

Therefore, 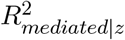 can be interpreted as the variation of the outcome variable that can be explained by the exposure variable through mediators adjusting for confounder(s) *Z*.

## 3. Results

In this section, we performed a series of simulation studies to assess the performance of the proposed MASH *R*^2^ mediation analysis procedure for time-to-event outcome in comparison with alternative procedures. We evaluated the MASH in two aspects—the estimated total mediation effect value and the true mediator selection and compared it with the state-of-art HIMA method [Luo et al. (2020)].

### 3.1. Simulation Design

We generated data using equations (2) and (4) . For subject *i* = 1, …, *n*, the exposure *X* was generated from the standard normal distribution *N* (0, 1), where the true mediators were generated using equation (4) and noise mediators were generated from *N* (0, 1). The error term *ϵ*_*ij*_ was generated from *N* (0, 1). We assumed a common form of baseline hazard *λ*_0_(*t*) = *λt*^*η−*1^ following a Weibull distribution with *λ* = 2 and *η* = *−*5. The censoring time was generated from *U* (0, *c*_0_) with a constant *c*_0_, which was chosen to control the censoring rate. Furthermore, we conducted simulation studies in 16 settings, varying the censoring rate, coefficients of mediators in true models, number of true mediators, and number of potential mediators. For each simulation setting, we generated 500 replicates, and the sample size *n* was 2000 to mimic our real data application to the FHS. To evaluate the performance of MASH in estimating the mediation effect, we calculated a pseudo-true *R*^2^ mediation effect with known parameters and true mediators in our simulation data-generation model. The pseudo-true *R*^2^ mediation effect is not the exact true value because the true parameters *c* and *d* are unknown, so the *R*^2^ of models (2) and (4) (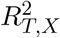 and 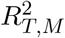) are estimated values, which may introduce bias and variance in model fitting. To circumvent the undue influence of model fitting uncertainty, we calculated the pseudo-true *R*^2^ mediation effect using a very large sample size (*n* = 200,000).

We performed simulation studies in each setting with censoring rates of 10%, 30%, and 50%. We found that the value of the estimated mediation effect increased with the censoring rate. Current censoring methods in simulation studies in the literature typically use a uniform or exponential distribution to generate the censoring time and adjust the parameters of these distributions to control the censoring rate. With a varying censoring rate, data will not have the same effective sample size. Therefore, comparing the results using censored data with those using uncensored data is unfair. Thus, we used a pseudo-true *R*^2^ mediation effect with comparable censoring rates (10%, 30%, and 50%). The bias for our estimated *R*^2^ mediation effect (shown in Fig. 3) increased very slightly as the censoring rate increased.

**FIG 3.**
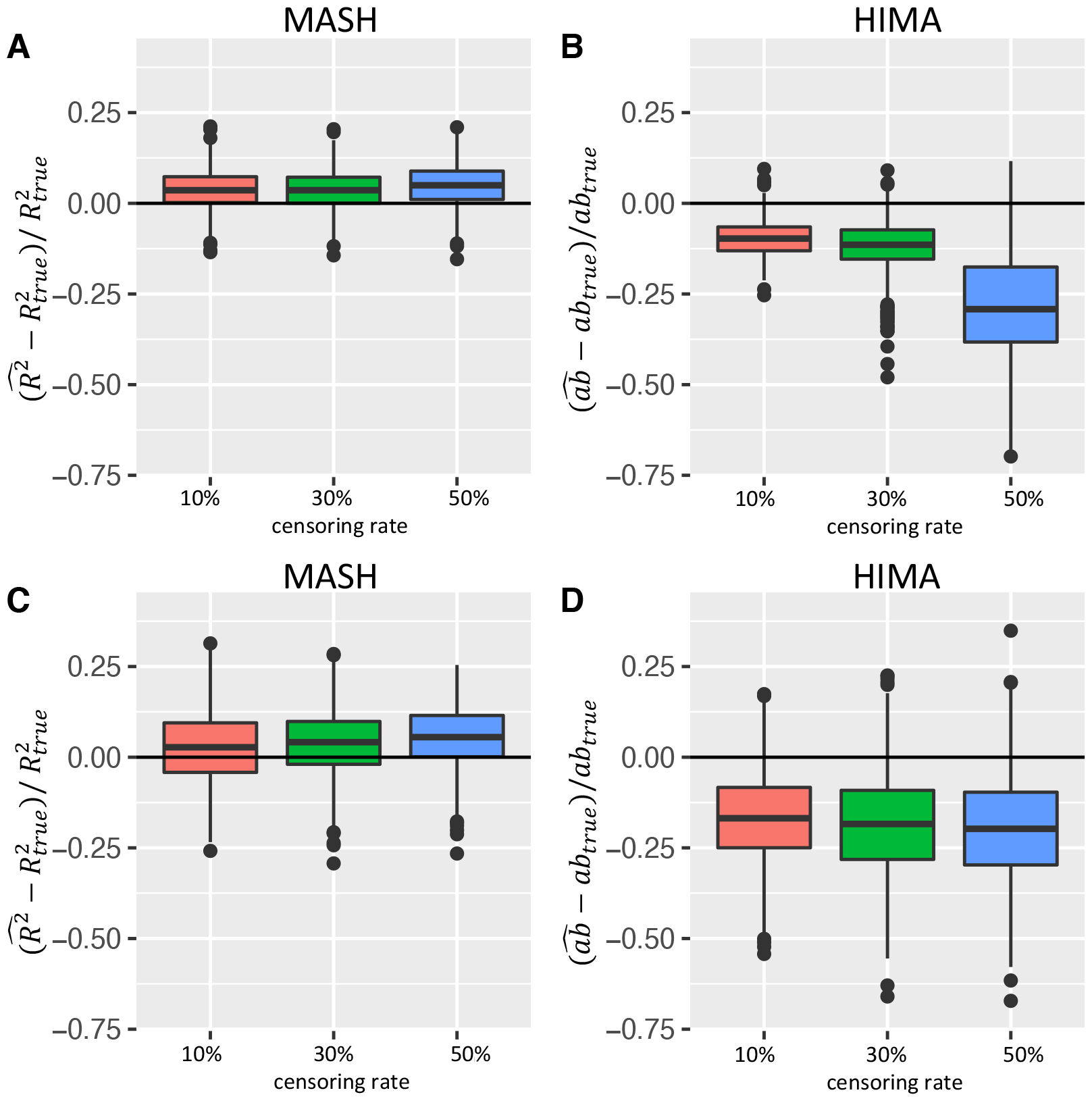
Boxplots of the relative bias across simulation replications of MASH and HIMA. (Simulation setting: *n* = 2000, *m* = 5, *p* = 1000; *a*_*j*_ = 0.4, *b*_*j*_ = 0.25 for *j* = 1, 2, …, 5 in setting 1 and *a*_*j*_ = 0.65 for *j* = 1, 2, …, 5, *b* = (-1, -1, 0.5, 0.5, 0.5) in setting 8.) The x-axis corresponds to the censoring rate at (10%, 30%, 50%); Y-axis corresponds relative bias across simulation replications. (A and B) mediation effects of all true mediators were in the same direction (setting 1); (C and D) mediation effects of true mediators were in different directions (setting 8). MASH results are shown in A and C, and HIMA results are shown in B and D.

### 3.2. Simulation results

To explore the performance of the MASH procedure in different scenarios, we conducted simulation studies in settings with different numbers of true mediators (*m* = 1, 5, and 10) and a fixed number of potential mediators (*p* = 1000), in settings with ultra-high-dimensional data (*p* = 1000, 5000, and 10,000, and *n* = 2000) and a fixed number of true mediators (*m* = 10), and in settings with different values of coefficients *a* and *b* when the numbers of true mediators *m* and potential mediators *p* were fixed. Simulation results for all settings are shown in the Supplementary Material Table S1-S7. The results demonstrated that our estimation of the mediation effect using MASH was very stable and performed well in terms of bias, variance and variable selection across the different numbers of true mediators *m* and a fixed *p* when we varied the values of parameters *a* and *b* and number of potential mediators while keeping *m* and *p* fixed. With ultra-high dimensional data (settings with *p* = 1000, 5000, and 10,000), MASH still performed well as shown in Tables S2, S3 and S6. With a small sample size (*n* = 300 and 500), the performance of our method could be biased as shown in Table S7. However, the bias is still very small if *n* = 500 and censoring rate is less or equal to 30%. Therefore, we would suggest using our method with reasonably large samples to avoid bias. More precisely, we need to consider the effective sample size for survival analysis, which is the number of non-censored subjects.

Figure 3 shows boxplots of the estimated total mediation effect of MASH and HIMA in two settings with *n* = 2000, *m* = 5, and *p* = 1000. The coefficients in the two settings were different. In setting 1, *a* = 0.4, and the coefficients of mediators *b* was 0.25 for all five true mediators. When the directions of component-wise mediation effects differ across multiple mediators, a potential problem is that the effects in different directions could cancel each other out in the product measure of the total mediation effect. Therefore, Figure 3 also shows results of setting 8 with five mediators that had different component-wise mediation effects directions with coefficients *b* = (-1, -1, 0.5, 0.5, 0.5). Figures 3A and 3C show MASH results, and Figure 3B and 3D shows HIMA results. In comparing MASH and HIMA in setting 1 (Fig. 3A and 3B), setting 8 (Fig. 3C and 3D) and all other settings described in the supplementary materials, we found that estimating the total mediation effect using MASH produced a smaller relative bias and smaller variance than did that using HIMA. Also, MASH was more stable than using the HIMA as the censoring rate increased as the product measure is only approximately true for a survival outcome in rare disease setting[VanderWeele et al. (2011)]. In addition, the bias in estimating *b* in Cox model increases as censoring rate increases. Therefore, the bias of the total indirect effect estimation in summation of *ab* further increases as the summation of censoring effects. However, *R*^2^ is an overall measure of explained variation using three Cox models, which is less dependent on individual parameter estimation accuracy and little affected by censoring [Schemper and Stare (1996)]. For setting 8, the total indirect effects using HIMA were negative, with the conflict directions of the component-wise mediation effects canceled out in the product measure when the total effect of exposure on outcome was positive. Overall, MASH outperformed HIMA in high-dimensional mediation analysis for survival outcomes across varying simulation settings.

To evaluate our proposed partial *R*^2^ method adjusting for confounders, we performed a simulation study using equations (9) and (11) as data-generating models. We used a sample size of 2000 and *p* = 1000 potential mediators, 5 of which were true mediators. We also used censoring rates of 10%, 30%, and 50% and *U* (0, 1) as the censoring time distribution. We set the parameters *a* = 0.38, *b* = 0.5, *r* = 2.5, and *l* = 0.1 for all true mediators. The mediator-outcome confounder *Z* and noise mediators were generated by the standard normal distribution. We compared our proposed partial *R*^2^ method with the unadjusted *R*^2^ method in this simulation setting, varying the coefficients of covariate *g* (0.5, 0.75, and 1). The simulation results shown in Supplementary Material Table S5 demonstrated that the partial *R*^2^ measure performed well in estimating the mediation effect controlling for confounding effects.

The existence of the non-mediators *U* and *V* in identified mediators set *S*_3_ could bias our estimation of the total mediation effect; thus, we considered two settings representing the scenarios that potential mediators set *S*_0_ included ten non-mediators *U* or *V* , the results of which are shown in the Supplementary Material Table S4. We used a sample size of 2000 and 1000 potential mediators, 1 of which was the true mediator.

We still used censoring rates of 10%, 30%, and 50% and *U* (0, 1) as the censoring time distribution. We also set the parameters *a* = 1, *b* = 0.5, and *r* = 2.5 for the true mediator. We set the parameters *a* = 1 and *b* = 0 for 10 non-mediators *V* and *a* = 0 and *b* = 0.5 for 10 non-mediators *U* . The results demonstrated that our proposed MASH consistently performs well in estimating the mediation effect, although the bias of the estimation slightly increased with non-mediators *U* and *V* presence.

In addition to the mediation effect estimation, we assessed the accuracy of mediator variable selection using MASH. The true positive rate (percentage of true mediators correctly selected) and false positive rate (percentage of non-mediators incorrectly selected) for all settings are presented in Supplementary Material Table S6. The true positive rates were greater than 99% among all settings, and the highest false positive rate for all settings was 0.7%. Therefore, the performance of MASH is satisfactory in terms of selecting true mediators and controlling the false positive rate.

## 4. Real Data Applications

To demonstrate the proposed MASH method’s versatility with different omics mediators and study designs, we applied it to the metabolomics data as mediators in the prospective cohort study FHS with time to coronary heart disease (CHD) and smoking-related cancer as the outcomes, as well as genome-wide copy-number variations as mediators with overall survival as the outcome in a cancer genomics dataset.

### 4.1. Application of MASH to the FHS

The FHS is a long-term cohort study of cardiovascular disease that has been ongoing since the 1940s and includes three generations: the Original Cohort, the Offspring Cohort, and the Third Generation Cohort [Splansky et al. (2007)]. In our application of MASH to the FHS, we used the metabolomics data of 1919 individuals in the Offspring Cohort from whom blood samples were collected at Exam 5. We were interested in exploring the roles of metabolites in the pathway from smoking to the risk of CHD or cancer.

#### 4.1.1. CHD CHD is the leading cause of death globally

Smoking is a major risk factor for CHD risk [HHS (2010)]. Although researchers have studied many metabolites and showed that they are significantly associated with the risk of incident CHD [e.g. Wang et al. (2019); Cavus et al. (2019); Li et al. (2017)], their mediation role in the smoking-CHD relationship has yet to be elucidated. Therefore, we applied MASH to the plasma-based metabolomics data from individuals in the FHS Offspring Cohort. After excluding subjects with more than 20% missing values in smoking status or metabolite data, we applied the k-nearest neighbors (KNN) algorithm [Altman (1992)] to impute the remaining missing data. After preprocessing, we had 1902 individuals, 307 of whom were diagnosed with CHD during follow-up after Exam 5 when plasma was collected for metabolomics profiling, with 190 metabolites as potential mediators. The exposure was tobacco smoking status at Exam 5 as a binary variable (never/ever). The outcome was the number of days from Exam 5 to CHD diagnosis or the last follow-up visit. The median number of follow-up days was 8672.5. We evaluated the effect of smoking on CHD risk as mediated by metabolites, adjusting for age, gender and body mass index (BMI). We randomly split the data into 50% as variable selection set and 50% as estimation set as illustrated in Fig. 2. We used the first half to select mediators and used the second half to estimate the mediation effect. The number of mediators *p* was smaller than the sample size *n* in the FHS data set, so we skipped the preliminary principled sure independence screening step here. We controlled false discovery rate at 20% in step (c). We also applied the HIMA method as comparison.

The results are presented in Table 1. We identified five metabolites as mediators of the effect of smoking on CHD risk including cotinine, carnitine, C48:0 triacylglycerol, C20:4 cholesteryl ester, and monosaccharides. The total mediation effect of the metabolites was 0.017 (SE = 0.016) by the 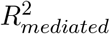measure. Because the values of 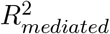 measures depend on the total effect, we focused on more interpretable *SOS* measures in real data applications, which are standardized by the total effect and can be interpreted as the percentage of the total effect that is explained by the mediators. We estimated that 51.1% (SE = 0.215) of the total effect was mediated by metabolites based on the *SOS* measure. We also evaluated the mediation effect of each selected mediators in single-mediator models and presented their mediation effects as estimated using the partial *R*^2^ method, adjusting for age, gender and BMI. In our single-mediator model, we estimated that cotinine, carnitine, C48:0 triacylglycerol, C20:4 cholesteryl ester, and monosaccharides explained 21.5%, 20.3%, 16.1%, 15.5%, and 9.6% of the total effect, respectively, by the *SOS* measure Cotinine is an alkaloid found in tobacco and is the predominant metabolite of nicotine. It is a highly sensitive and specific marker of active and passive exposure to tobacco [Benowitz (1996)]. Researchers reported similarity in the relative risk of CHD among individuals with substantial exposure to passive smoking and those with light active exposure to it [Whincup et al. (2004)]. Also, studies have demonstrated that the ability of urine cotinine to improve assessment of cardiovascular disease risk assessment is similar to that provided by self-reported smoking status [Kunutsor et al. (2018)]. Our results further showed that cotinine is a mediator in the pathway from smoking to the risk of CHD.

**TABLE 1.**
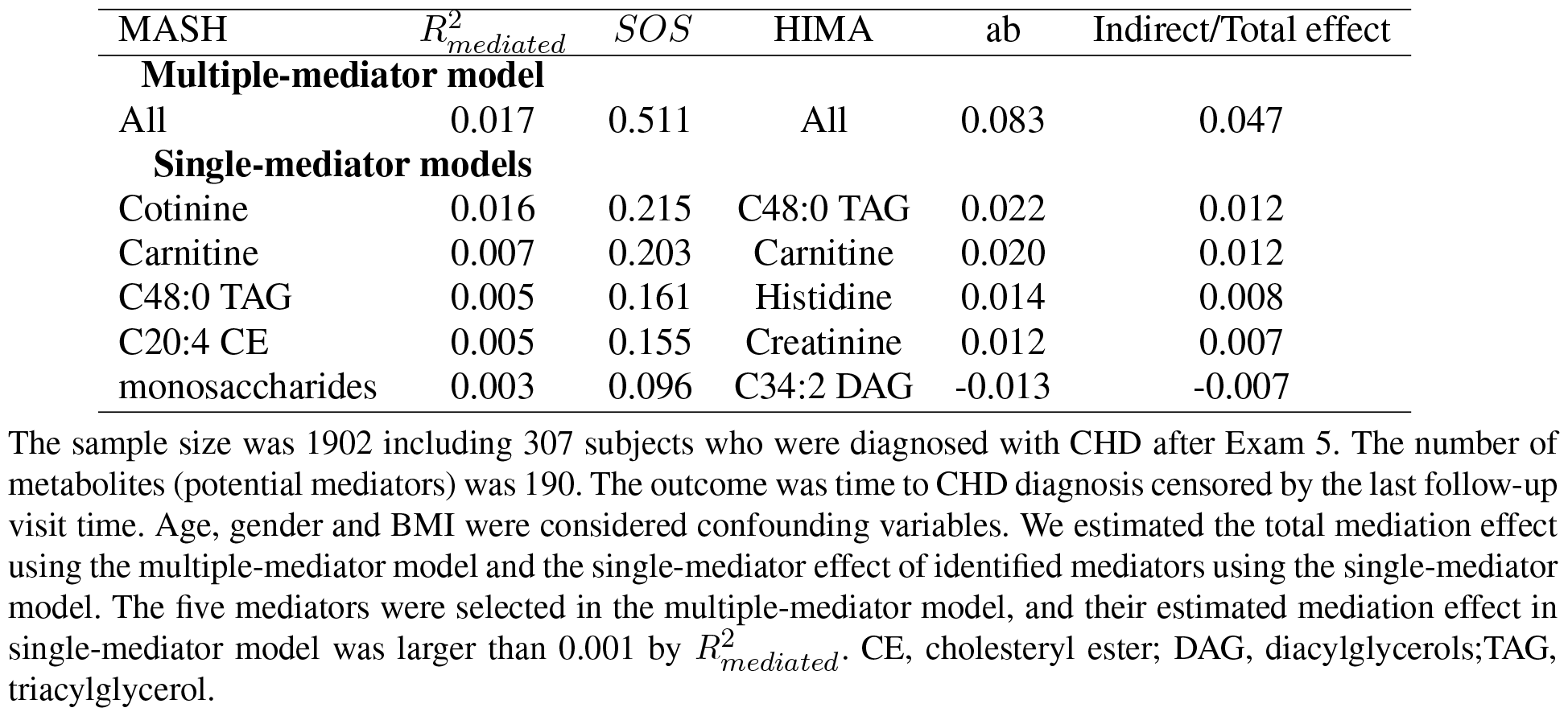
Application of proposed MASH to FHS data: smoking, metabolites and CHD risk in comparison with HIMA.

| MASH | $R^2_{mediated}$ | SOS | HIMA | ab | Indirect/Total effect |
| --- | --- | --- | --- | --- | --- |
| **Multiple-mediator model** |  |  |  |  |  |
| All | 0.017 | 0.511 | All | 0.083 | 0.047 |
| **Single-mediator models** |  |  |  |  |  |
| Cotinine | 0.016 | 0.215 | C48:0 TAG | 0.022 | 0.012 |
| Carnitine | 0.007 | 0.203 | Carnitine | 0.020 | 0.012 |
| C48:0 TAG | 0.005 | 0.161 | Histidine | 0.014 | 0.008 |
| C20:4 CE | 0.005 | 0.155 | Creatinine | 0.012 | 0.007 |
| monosaccharides | 0.003 | 0.096 | C34:2 DAG | -0.013 | -0.007 |

The sample size was 1902 including 307 subjects who were diagnosed with CHD after Exam 5. The number of metabolites (potential mediators) was 190. The outcome was time to CHD diagnosis censored by the last follow-up visit time. Age, gender and BMI were considered confounding variables. We estimated the total mediation effect using the multiple-mediator model and the single-mediator effect of identified mediators using the single-mediator model. The five mediators were selected in the multiple-mediator model, and their estimated mediation effect in single-mediator model was larger than 0.001 by $R^2_{mediated}$ . CE, cholesteryl ester; DAG, diacylglycerols; TAG, triacylglycerol.

Previous studies also found that L-carnitine can effectively improve the cardiac function rating and thus improve cardiac function [Zhao et al. (2020)] and was associated with a 27% reduction in the all-cause mortality rate, a 65% reduction in ventricular arrhythmias, and a 40% reduction in anginal symptoms in patients experiencing acute myocardial infarction [DiNicolantonio et al. (2013)]. Carnitine is also a metabolite found at higher levels in current smokers than in non-smokers [Xu et al. (2013)]. Our results are consistent with the results of these studies and revealed the important mediation role of carnitine in the impact of smoking on the risk of CHD.

Cholesteryl ester present in human plasma is positively correlated with high-density lipoprotein (HDL) levels [Subbaiah et al. (2012)]. Many clinical and epidemiological studies have clearly demonstrated that the HDL cholesterol level is inversely associated with the risk of CHD and is a critical and independent component of predicting this risk [Wang et al. (2017); Mundra et al. (2018)]. In addition, elevated plasma triacylglycerol concentrations have been associated with increased risk of CHD and associated with other CHD risk factors, namely, reduced HDL cholesterol concentrations and increased low-density lipoprotein (LDL) particles [Roche and Gibney (2000)]. Previous studies suggested that smoking is associated with total cholesterol, low-density lipoprotein cholesterol, and triglyceride levels [Gossett et al. (2009)]. Our mediation analysis results provide further understanding of how these risk factors work together to contribute to the incidence of CHD.

The three most common monosaccharides are glucose, fructose, and galactose. Sugars are naturally components of the tobacco leaf and are also commonly added to cigarettes by tobacco companies. Glucose and fructose are the most abundant natural sugars in dried tobacco leaves Clarke et al. (2006). Many studies found that increased intake of sugar and blood sugar level were associated with an increased CHD risk Howard et al. (2002). We confirmed the mediating role of monosaccharides in the effect of cigarette smoking on CHD risk using MASH.

As shown in Table 1, the indirect effect estimated using HIMA was 0.083, and the ratio of the indirect effect to the total effect was 0.047. HIMA selected 14 mediators, 3 of which were selected by MASH: carnitine, C48:0 TAG and monosaccharides. The 11 other mediators selected by HIMA were histidine, creatinine, adma, aminoadipate, amp, salicylurate, C36:1pc, C16:1sm, C34:2dag, C52:3tag, and C58:12tag. We observed conflicting directions of the indirect effect in product measure, which caused cancellation of the estimated total indirect effect by HIMA.

#### 4.1.2. Smoking-Related Cancer

Smoking can cause cancer and, further, prevent the body from fighting it [HHS (2020a)]. Specifically, it can cause cancer at almost all of the most common sites, including the bladder, colon, kidney, lung, and pancreas [HHS (2020b)]. Many studies have suggested that metabolites are important biomarkers associated with the risk of cancer [Mazzilli et al. (2020)]. In addition, smoking is associated with many metabolites, such as cotinine, O-cresol sulfate and hydroxycotinine [Cross et al. (2014)]. The risk of colon, lung, pancreatic, urinary bladder, and kidney cancer was shown to be associated with cigarette smoking in many previous studies [ACS (2021); CDC (2020)]. Therefore, the mediation role of metabolites in the relationship between tobacco smoking and cancer is of great interest. To examine this, we applied the MASH method to the FHS Offspring Cohort with smoking as the exposure, metabolomics at Exam 5 as the potential mediators and incident smoking-related cancer during the follow-up after Exam 5 as the outcome, adjusting for age, gender and BMI as covariates. We were left with 1919 individuals and 190 metabolites after removing subjects and metabolites with more than 20% missing values. One hundred eighty-five individuals in the data set were diagnosed with the aforementioned smoking-related cancer during follow-up. As illustrated in Figure 2, we randomly spit the data set into two halves for variable selection and estimation, respectively.

The results are presented in Table 2. We identified two metabolites that mediated the effect of smoking on the risk of cancer: cotinine and glycodeoxycholic acid. The total mediation effect of the metabolites was 0.055 (SE = 0.026) by the 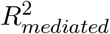 measure. We estimated that 50.7% (SE = 0.184) of the total effect was mediated by the two metabolites according to the *SOS* measure. In our single-mediator models, we estimated that cotinine explained 46.8% of the total effect, and glycodeoxycholic acid explained 4.2% of the total effect by the *SOS* measure. Previous studies demonstrated that cotinine, as a biomarker of current smoking status, was associated with the incidence of bladder cancer [Thong et al. (2016)], lung cancer [Ellard et al. (1995)], colorectal cancer [Cross et al. (2014)], and other smoking-related cancers. Furthermore, it was reported that glycodeoxycholic acid, a conjugated bile acid, stimulated tumor growth [Dai et al. (2013)], and a recent study suggested that it promotes colon cancer development [Kühn et al. (2020)]. On the other hand, HIMA selected 0 mediator, which means that the estimated mediation effects of metabolites in the pathway from smoking to cancer using HIMA was 0. Therefore, MASH produced more reasonable results of mediation effect estimation than HIMA in the applications of real data with high censoring rates.

**TABLE 2.** Application of proposed MASH to FHS data: smoking, metabolites and cancer risk.

| Mediators | $R^2_{mediated}$ | *SOS* |
| --- | --- | --- |
| **Multiple-mediator model** |  |  |
| All | 0.055 | 0.507 |
| **Single-mediator model** |  |  |
| Cotinine | 0.051 | 0.468 |
| Glycodeoxycholic | 0.005 | 0.042 |

The sample size was 1919 including 185 individuals who were diagnosed with smoking-related cancer after Exam 5. The number of metabolites (potential mediators) was 190. The outcome was time to cancer diagnosis censored by the last follow-up visit time. Age, gender and BMI were considered as confounding variables. We estimated total mediation effect using the multiple-mediator model and the single-mediator effect of identified mediators using the single-mediator model. The two mediators were selected in the multiple-mediator model, and their estimated mediation effect in the single-mediator model was larger than 0.001 by $R^2_{mediated}$ .

### 4.2. Prognosis for DLBCL

Diffuse large B-cell lymphoma (DLBCL) is the most common hematological malignancy and is characterized by a striking degree of genetic and clinical heterogeneity [Reddy et al. (2017); Xu et al. (2022)]. The International Prognostic Index (IPI), a clinical scoring system developed by oncologists to aid in predicting the prognosis for lymphoma, assigns one point to each negative prognostic factor (age > 60 years, serum lactate dehydrogenase level above the upper limit of normal, Ann Arbor stage III/IV disease, Eastern Cooperative Oncology Group performance status 2, and more than one site with extranodal involvement) and categorizes patients into four risk groups based on the total score: 0/1 = low risk, 2 = low-intermediate risk, 3 = high-intermediate risk, and 4/5 = high risk. Researchers developed the IPI at a time when DLBCL patients received chemotherapy-only regimens, and had estimated 5-year overall survival rates ranging from 26% to 73% depending on the risk category [INHLPFP (1993)]. We were interested in determining how the clinical exposure (IPI score) affects the overall survival of DLBCL patients through copy-number alterations. Reddy et al. (2017) published a study of 1001 DLBCL patients with copy-number alteration profiling of 140 genes known to be associated with cancer development and prognosis. We used 754 observations in this data set for the present study, after removing 247 patients with missing IPI scores or outcomes (overall survival in years). The censoring rate was 0.683. In our application of MASH, we randomly spit the data set into two halves for variable selection and estimation. Our results shown in Table 3 demonstrated that the total mediation effect of the copy-number alterations was 0.063 (SE = 0.041) by the 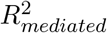 measure. And we estimated that the total mediation effect of the copy-number alterations explained 19.7% (SE = 0.130) of the total effect by the *SOS* measure. We identified eight genes that mediated the effect of IPI score on the overall survival using our MASH procedure. In our single-mediator models, we estimated that *FOXP*1, *POU*2*F* 2, *ANKRD*17, *LIN* 54, *CD*70, *DNMT* 3*A, S*1*PR*2, and *CD*79*B* explained 8.3%, 7.4%, 6.0%, 4.0%, 3.3%, 1.9%, 0.6% and 0.6% of the total effect by the *SOS* measure, respectively. In another study, investigators identified *ANKRD*17 [Reddy et al. (2017)], *POU* 2*F* 2 [Reddy et al. (2017)], *CD*79*B* [Reddy et al. (2017)], and *S*1*PR*2 [Baldari et al. (2016)] as driver genes for DLBCL. Researchers also found that the genes *DNMT* 3*A* [Reddy et al. (2017)], *CD*70 [Reddy et al. (2017)], *CD*79*B* [Schmitz et al. (2018)], and *FOXP* 1 [Barrans et al. (2004)] are prognostic factors for DLBCL overall survival. *LIN*54, a protein coding gene, is a component of the *LIN* , or *DREAM* complex, which is an essential regulator of cell cycle genes known to be associated with ovarian cancer. While looking into the gene expression and copy number variation of *LIN*54, we found that they were both borderline significant for overall survival of BLDCL patients (Supplemental materials Section 4).

**TABLE 3.** Application of proposed MASH to DLBCL prognosis: IPI score, copy-number alterations and overall survival of DLBCL patients.

| Mediators | $R^2_{mediated}$ | SOS |
| --- | --- | --- |
| **Multiple-mediator model** |  |  |
| All | 0.063 | 0.197 |
| **Single-mediator model** |  |  |
| FOXP1 | 0.027 | 0.083 |
| POU2F2 | 0.024 | 0.074 |
| ANKRD17 | 0.019 | 0.060 |
| LIN54 | 0.013 | 0.040 |
| CD70 | 0.011 | 0.033 |
| DNMT3A | 0.006 | 0.019 |
| S1PR2 | 0.002 | 0.006 |
| CD79B | 0.002 | 0.006 |

The sample size was 754 and censoring rate was 68.3%. The number of genes of copy-number alterations (potential mediators) for each patient was 140. The outcome was overall survival. We estimated total mediation effect using the multiple-mediator model and the single-mediator effect of identified mediators using the single-mediator model. The eight mediators were selected in the multiple-mediator model, and their estimated mediation effect in the single-mediator model was larger than 0.001.

Lastly, HIMA selected 0 mediator, which means that the estimated mediation effect of copy number variation in the pathway from clinical IPI score to DLBCL using HIMA was 0. Therefore, MASH produced more reasonable results of mediation effect estimation than HIMA and outperformed HIMA in the three applications of real data with high censoring rates.

## 5. Discussion

We have developed MASH, a novel method of mediation analysis, for high-dimensional mediators and time-to-event outcomes to estimate the total mediation effect and identify mediators. MASH critically extends the existing *R*^2^-based mediation analysis for continuous outcomes to survival outcomes in the high-dimensional setting. Our approach incorporates sure independence screening, MCP variable selection, and false discovery rate control for high-dimensional mediator variable selection and *R*^2^-based measures for mediation effect estimation. It has good performance in mediation effect estimation and mediator selection, which we have shown via simulation studies and multiple real data applications. Extending the *R*^2^ measure defined in linear regression to survival models is not straight-forward owing to the nonunique definition of *R*^2^ in the latter. Based on our simulations, the 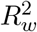 -based measure should be chosen for survival models because the influence of censoring on it is less than that on other *R*^2^ measures. Besides the Cox proportional hazards model, we have also explored an accelerated failure time (AFT) model for analysis of right censored data and found that the Cox model is more stable in *R*^2^ measure estimation with a wide range of censoring rates.

Many questions of interest about mediation analysis for high-dimensional survival data remain to be addressed in the future. Mediation analysis of multiple types of high-dimensional omics mediators such as gene expression data in addition to metabolomics data, is biologically interesting, yet challenging for selecting mediators from multi-omics data with potentially complex dependence structures. Although we have proposed using partial *R*^2^ measure for MASH to control confounding in mediation analysis, models allowing for exposure-mediator interactions warrant further investigation in the future.

## Acknowledgements

This work was supported by the National Institutes of Health (NIH) grants R01HL116720. P.W. was partially supported by NIH grant P50CA217674. X.H. was partially supported by NIH grants R01CA272806, U54CA096300, U01CA152958 and P50CA100632, and the Dr. Mien-Chie Hung and Mrs. Kinglan Hung Endowed Professorship. The authors acknowledge the Texas Advanced Computing Center at The University of Texas at Austin for providing HPC resources. The authors declare that there are no conflicts of interest. The Framingham Heart Study is conducted and supported by the National Heart, Lung, and Blood Institute (NHLBI) in collaboration with Boston University (Contract No. N01-HC-25195). This article was not prepared in collaboration with investigators of the Framingham Heart Study and does not necessarily reflect the opinions or views of the Framingham Heart Study, Boston University, or NHLBI. The manuscript was edited by Don Norwood and Sarah Bronson, ELS, of Editing Services, Research Medical Library at The University of Texas MD Anderson Cancer Center.

## SUPPLEMENTARY MATERIAL

**Supplement to “MASH: Mediation analysis of survival outcome and high-dimensional omics mediators with application to complex disease.”**

The supplementary material contains the complete results of our simulation study discussed in the text.

