## Supplemental materials for "MASH: Mediation Analysis of Survival Outcome and High-dimensional Omics Mediators with Application to Complex Diseases"

#### 1 Derivation of $R^2_{mediated}$ in the Cox proportional hazards model

Here we present the calculation of $R^2_{mediated}$ using coefficients and variance matrix under models:

$$\lambda(t|X_i) = \lambda_0^X(t)exp(cX_i), \tag{1}$$

$$\lambda(t|X_i, M_i) = \lambda_0^{XM}(t)exp(rX_i + \sum_{j=1}^{p} b_j M_{ij}), \tag{2}$$

$$\lambda(t|M_i) = \lambda_0^M(t)exp(\sum_{j=1}^{p} d_j M_{ij}), \tag{3}$$

$$M_{ij} = a_j X_i + \epsilon_{ij}, \tag{4}$$

where $M_i = (M_{i1}, M_{i2}, \ldots, M_{ip})'$ is the $p$-dimensional mediator vector for subject $i = 1, 2, 3, ..., n$ and, without loss of generality, $X$ and $M_j$ are standardized to have mean 0 and variance 1. Let $D$ denote the time to event and $C$ denote the censoring time. The observed survival outcome is $T_i = min(D_i, C_i)$, and the failure indicator is $\delta_i = I(D_i \leq C_i)$ for $i = 1, 2, 3, ..., n$. Equations (1), (2), and (3) are Cox proportional hazards models describing the relationship (1) between $T$ and $X$, (2) between $T$, $X$, and $M$, and (3) between $T$ and $M$, respectively. $\lambda_0^X(t), \lambda_0^{XM}(t)$, and $\lambda_0^M(t)$ are their respective baseline hazard functions. In a Cox proportional hazards regression model, the hazard ratio is usually used as a measure of an independent variable's effect on a dependent variable. In equations (1), (2), and (3), $c$ is the parameter relating the exposure to the outcome; $r$ is the direct effect parameter relating the exposure to the outcome; $b$, defined as $(b_1, \ldots, b_p)^T$, is the parameter vector relating the mediators to the outcome with adjustment for the effect of the exposure; and $d = (d_1, \ldots, d_p)'$, is the parameter vector relating the mediators to the outcome. Equation (4) characterizes how exposure influences the mediators, where $a = (a_1, \ldots, a_p)'$ is the parameter vector relating the exposure to the mediators and residual $\epsilon_i = (\epsilon_{i1}, \epsilon_{i2}, \ldots, \epsilon_{ip}) \sim MVN(0, E_{p\times p})$, for $i = 1, \ldots, n$.

$R^2$ [1] of model (1) is

$$R^2_{T,X} = \frac{var(cX)}{1 + var(cX)} = \frac{c^2}{1 + c^2}, \tag{5}$$

$R^2$ of model (2) is

$$R^2_{T,MX} = \frac{var(rX + \sum_{j=1}^{p} b_j M_j)}{1 + var(rX + \sum_{j=1}^{p} b_j M_j)} = \frac{(r + b^T a)^2 + b^T Eb}{1 + (r + b^T a)^2 + b^T Eb}, \tag{6}$$

$R^2$ of model (3) is

$$R^2_{T,M} = \frac{var(\sum_{j=1}^{p} d_j M_j)}{1 + var(\sum_{j=1}^{p} d_j M_j)} = \frac{(d^2)^T (a^2) + d^T Ed}{1 + (d^2)^T (a^2) + d^T Ed}, \tag{7}$$

such that

$$R^2_{mediated} = R^2_{T,M} + R^2_{T,X} - R^2_{T,MX} = \frac{(d^2)^T (a^2) + d^T Ed}{1 + (d^2)^T (a^2) + d^T Ed} + \frac{c^2}{1 + c^2} - \frac{(r + b^T a)^2 + b^T Eb}{1 + (r + b^T a)^2 + b^T Eb}, \tag{8}$$

#### 2 Relationship between the product measure and $R^2_{mediated}$

Under the classical causal mediation analysis framework, the standard indirect effect measure, product measure, is defined by $b^T a$. Therefore, a potential mediator with either $a$ or $b$ equal to 0 will be identified as non-mediator. To investigate the relationship between the product measure and $R^2_{mediated}$ in the scenarios with no indirect effect by the product measure, i.e., $b^T a = 0$.
In the scenario of all $a$'s and $b$'s equal to 0, $c = r$ and $d = b = 0$,

$$R^2_{mediated} = \frac{d^T Ed}{1 + d^T Ed} + \frac{c^2}{1 + c^2} - \frac{r^2}{1 + r^2} = 0. \tag{9}$$

Therefore, both $R^2_{mediated}$ and product measure demonstrate no mediation effect if all $a$'s and all $b$'s equal to 0. We examined the performance of MASH in this scenario in comparison with HIMA. The mean and variance (in parentheses) of $R^2_{mediated}$, SOS and HIMA are presented in the Table S1. The results showed that both our approach MASH and HIMA are essentially unbiased when all $a$'s and $b$'s are zero. Given the relationship between $R^2_{mediated}$ and the product measure, even when none of $a_j$'s or $b_j$'s is zero, the product measure $b^T a$ could be zero or close to zero due to cancellation of positive and negative $a_j b_j$'s, while $R^2_{mediated}$ can be away from zero. Then we present the U-type and V-type mediators when $R^2_{mediated}$ is misleadingly non-zero and product measure is correctly zero. Therefore, we need the variable selection procedures in MASH. Please refer to Table S4 for further simulation studies with U-type or V-type mediators (only $a_j$'s or $b_j$'s equal to 0).
In the scenario of all $a_j$'s equal to 0 but none of $b_j$'s equal to 0, it follows that variables in $M$ are all U-type non-mediators (Figure 1C in the main text). Since that all $a_j$'s equal to 0 implies that $M \perp\!\!\!\perp X$, $c = r$ and $b = d$, it follows that

$$R^2_{mediated} = \frac{d^T Ed}{1 + d^T Ed} + \frac{c^2}{1 + c^2} - \frac{r^2 + b^T Eb}{1 + r^2 + b^T Eb} = \frac{d^T Ed}{1 + d^T Ed} + \frac{r^2}{1 + r^2} - \frac{r^2 + d^T Ed}{1 + r^2 + d^T Ed}, \tag{10}$$

so $R^2_{mediated}$ is typically not zero under this scenario.
In the scenario of all $b_j$'s equal to 0 but none of $a_j$'s is equal to 0, $c = r$ and $M \perp\!\!\!\perp Y|X$, which indicates $M_j$'s are V-type non-mediators as in Figure 1D in the main text.

$$R^2_{mediated} = \frac{(d^2)^T a^2 + d^T Ed}{1 + (d^2)^T a^2 + d^T Ed} + \frac{c^2}{1 + c^2} - \frac{r^2}{1 + r^2} = \frac{(d^2)^T a^2 + d^T Ed}{1 + (d^2)^T a^2 + d^T Ed}, \tag{11}$$

Table S1: Simulation results when all $a$'s and $b$'s are zero (i.e., both $R^2_{mediated}$ and the product measure are zero)

| Censoring | $R^2_{mediated}$ | SOS | HIMA |
|---|---|---|---|
| 10% | 0.004 | 0.005 | <-0.001 |
| | (0.005) | (0.005) | (0.002) |
| 30% | 0.005 | 0.005 | -0.002 |
| | (0.005) | (0.006) | (0.004) |
| 50% | 0.006 | 0.007 | 0.001 |
| | (0.007) | (0.008) | (0.005) |

#### 3 Simulation studies

We performed a series of simulation studies to assess the performance of our proposed MASH procedure in two aspects——the estimated total mediation effect value and true mediator identification. We use seven supplementary tables to present results of 17 simulation settings we conducted, where we evaluate our total mediation effect estimation performance in terms of the bias and variance. (H1) to (H8) in Table S1 and S2 represent the scenarios of high-dimensional settings with fixed sample size $n = 2000$ and number of potential mediators $p = 1000$, but varying the censoring rate, coefficients of mediators, and number of true mediators. In (H8), we let the true mediators have different directions of mediation effects. (H$_{mimic}$) in Table S1 represent the scenarios mimicking the real data. In Table

S3, (H9), (H10) and (H11) represent ultra-high dimensional scenarios. And we evaluated the effect of non-mediators U and V under (H12) and (H13) shown in Table S4. The situations that confounding occurred were represented by (H14), (H15) and (H16) in Table S5 and S7. Table S6 shows the results on true mediator identification in all 16 settings in terms of true positive rate and false positive rate. Table S7 shows the results of high dimensional settings with small to moderate sample sizes.

Table S2: Bias and standard deviation under high-dimensional settings

| | m | Censoring | $R^2_{mediated}$ | $\widehat{R^2}_{mediated} - R^2_{mediated}$ | $ab$ | $\widehat{ab} - ab$ | $\widehat{ab}_{HIMA} - ab$ |
|---|---|---|---|---|---|---|---|
| H1 | 1 | 10% | 0.495 | 0.014 | 0.451 | 0.206 | 0.202 |
| | | | | (0.028) | | (0.038) | (0.026) |
| | | 30% | 0.511 | 0.016 | 0.444 | 0.221 | 0.226 |
| | | | | (0.028) | | (0.042) | (0.039) |
| | | 50% | 0.492 | 0.037 | 0.433 | 0.255 | 0.265 |
| | | | | (0.031) | | (0.050) | (0.081) |
| H2 | 5 | 10% | 0.486 | 0.038 | 0.339 | 0.180 | 0.190 |
| | | | | (0.025) | | (0.038) | (0.029) |
| | | 30% | 0.509 | 0.006 | 0.332 | 0.226 | 0.251 |
| | | | | (0.028) | | (0.042) | (0.027) |
| | | 50% | 0.530 | 0.040 | 0.306 | 0.266 | 0.374 |
| | | | | (0.030) | | (0.047) | (0.054) |
| H3 | 5 | 10% | 0.498 | 0.015 | 0.385 | 0.003 | 0.001 |
| | | | | (0.026) | | (0.038) | (0.027) |
| | | 30% | 0.519 | 0.019 | 0.380 | 0.001 | -0.029 |
| | | | | (0.027) | | (0.042) | (0.056) |
| | | 50% | 0.538 | 0.023 | 0.372 | 0.003 | -0.126 |
| | | | | (0.029) | | (0.049) | (0.079) |
| H4 | 5 | 10% | 0.483 | 0.017 | 0.515 | 0.006 | -0.415 |
| | | | | (0.029) | | (0.039) | (0.027) |
| | | 30% | 0.506 | 0.018 | 0.510 | 0.005 | -0.413 |
| | | | | (0.028) | | (0.043) | (0.029) |
| | | 50% | 0.529 | 0.025 | 0.496 | 0.003 | -0.406 |
| | | | | (0.031) | | (0.049) | (0.072) |
| H5 | 10 | 10% | 0.491 | 0.019 | 0.513 | 0.004 | -0.468 |
| | | | | (0.026) | | (0.040) | (0.028) |
| | | 30% | 0.504 | 0.025 | 0.500 | 0.006 | -0.458 |
| | | | | (0.028) | | (0.043) | (0.057) |
| | | 50% | 0.535 | 0.029 | 0.494 | 0.009 | -0.456 |
| | | | | (0.029) | | (0.050) | (0.081) |
| H6 | 10 | 10% | 0.480 | 0.020 | 0.596 | 0.005 | -0.539 |
| | | | | (0.027) | | (0.041) | (0.028) |
| | | 30% | 0.494 | 0.021 | 0.582 | -0.001 | -0.529 |
| | | | | (0.028) | | (0.043) | (0.042) |
| | | 50% | 0.530 | 0.029 | 0.577 | 0.009 | -0.529 |
| | | | | (0.027) | | (0.049) | (0.089) |
| H7 | 10 | 10% | 0.471 | 0.020 | 0.672 | 0.004 | -0.606 |
| | | | | (0.026) | | (0.044) | (0.030) |
| | | 30% | 0.494 | 0.023 | 0.661 | 0.002 | -0.597 |
| | | | | (0.028) | | (0.047) | (0.034) |
| | | 50% | 0.526 | 0.028 | 0.653 | 0.005 | -0.594 |
| | | | | (0.027) | | (0.050) | (0.079) |
| H8 | 5 | 10% | 0.321 | 0.009 | -0.265 | -0.007 | -0.005 |
| | | | | (0.032) | | (0.063) | (0.040) |
| | | 30% | 0.352 | 0.013 | -0.261 | -0.006 | -0.004 |
| | | | | (0.032) | | (0.070) | (0.047) |
| | | 50% | 0.383 | 0.021 | -0.254 | -0.008 | -0.007 |
| | | | | (0.033) | | (0.075) | (0.052) |
| $H_{mimic}$ | 10 | 50% | 0.579 | 0.008 | 0.532 | 0.002 | -0.069 |
| | | | | (0.029) | | (0.049) | (0.070) |
| | | 70% | 0.621 | 0.008 | 0.532 | -0.001 | -0.193 |
| | | | | (0.030) | | (0.064) | (0.096) |
| | | 90% | 0.673 | -0.013 | 0.530 | -0.045 | -0.316 |
| | | | | (0.069) | | (0.127) | (0.085) |

Simulation settings (H1)-(H8): sample size $n = 2000$; number of potential mediators $p = 1000$; coefficient $r = 2.5$; coefficients $a = (1, 0.4, 0.43, 0.38, 0.275, 0.256, 0.24, 0.65)$ for settings (H1)-(H8), respectively; coefficients $b = (0.5, 0.25, 0.2, 0.3, 0.2, 0.25, 0.3)$ for settings (H1)-(H7), respectively. In (H8), coefficients for the 5 true mediators $b = (-1,-1,0.5,0.5,0.5)$. In ($H_{mimic}$), we mimic the real data: $n = 2000$; $p = 200$; $r = 2.5$; $a = 0.275$; $b = 0.2$. Abbreviations: m, number of true mediators; $R^2_{mediated}$, true value of $R^2_{mediated}$; $\widehat{R^2}_{mediated}$, estimation of $R^2_{mediated}$ by MASH; $ab$ true value of product measure; $\widehat{ab}$, estimation of $ab$ by MASH; $\widehat{ab}_{HIMA}$, estimation of $ab$ by HIMA. For estimations, the first row is the bias and the second row is the corresponding standard deviation.

Table S3: Bias and standard deviation under ultra high-dimensional settings

| | p | Censoring | $R^2_{mediated}$ | $\widehat{R^2}_{mediated} - R^2_{mediated}$ |
|---|---|---|---|---|
| H9 | 2000 | 10% | 0.492 | 0.020 |
| | | | | (0.025) |
| | | 30% | 0.515 | 0.023 |
| | | | | (0.028) |
| | | 50% | 0.537 | 0.029 |
| | | | | (0.028) |
| H10 | 5000 | 10% | 0.492 | 0.029 |
| | | | | (0.026) |
| | | 30% | 0.515 | 0.031 |
| | | | | (0.027) |
| | | 50% | 0.537 | 0.025 |
| | | | | (0.035) |
| H11 | 10000 | 10% | 0.492 | 0.035 |
| | | | | (0.027) |
| | | 30% | 0.515 | 0.021 |
| | | | | (0.032) |
| | | 50% | 0.537 | -0.027 |
| | | | | (0.059) |

Simulation settings (H9)-(H11): $n = 2000$, $m = 10$, coefficient $a = 0.275$, $b = 0.2$, and $r = 2.5$. Abbreviations: p, number of potential mediators; $R^2_{mediated}$, true value of $R^2_{mediated}$; $\widehat{R^2}_{mediated}$, estimation of $R^2_{mediated}$ by MASH. For $\widehat{R^2}_{mediated} - R^2_{mediated}$, the first row is the bias and the second row is the corresponding standard deviation in brackets.

Table S4: Results of simulation studies using data with all types of non-mediators

| | | Censoring | $R^2_{mediated}$ | $\widehat{R^2}_{mediated} - R^2_{mediated}$ (not screening non-mediators) | $\widehat{R^2}_{mediated} - R^2_{mediated}$ (screening out non-mediators) |
|---|---|---|---|---|---|
| H12 | U | 10% | 0.471 | 0.353 | -0.017 |
| | | | | (0.007) | (0.027) |
| | | 30% | 0.494 | 0.340 | -0.011 |
| | | | | (0.006) | (0.026) |
| | | 50% | 0.526 | 0.312 | -0.004 |
| | | | | (0.008) | (0.029) |
| H13 | V | 10% | 0.471 | 0.286 | 0.044 |
| | | | | (0.010) | (0.041) |
| | | 30% | 0.494 | 0.277 | 0.049 |
| | | | | (0.012) | (0.041) |
| | | 50% | 0.526 | 0.256 | 0.053 |
| | | | | (0.013) | (0.040) |

Simulation settings (H12) and (H13): $n = 2000$, $p = 1000$, $m = 10$, coefficient $a = 0.24$, $b = 0.3$, and $r = 2.5$. For (H12), there are 10 U-type non-mediators with $a = 0$, $b = 0.3$. For (H13), there are 10 V-type non-mediators with $a = 0.24$, $b = 0$. Abbreviations: U, non-mediators associated with the outcome but not the exposure; V, non-mediators associated with the exposure but not associated with the outcome; $R^2_{mediated}$, true value of $R^2_{mediated}$; $\widehat{R^2}_{mediated}$ (not screening non-mediators), estimation of $R^2_{mediated}$ using all potential mediators; $\widehat{R^2}_{mediated}$ (screening out non-mediators), estimation of $R^2_{mediated}$ by MASH using our proposed mediator selection procedure to filter out non-mediators. For $\widehat{R^2}_{mediated} - R^2_{mediated}$, the first row is the bias and the second row is the corresponding standard deviation in brackets.

Table S5: Results of simulation studies using data with a confounder under $R^2$-like measure

| | g | Censoring | $R^2_{mediated}$ | partial$\widehat{R^2}_{mediated}$ | unadjust$\widehat{R^2}_{mediated}$ | $SOS$ | partial$\widehat{SOS}$ | unadjust$\widehat{SOS}$ |
|---|---|---|---|---|---|---|---|---|
| H14 | 0.5 | 10% | 0.548 | 0.012 | 0.027 | 0.656 | 0.012 | 0.026 |
| | | | | (0.025) | (0.024) | | (0.026) | (0.025) |
| | | 30% | 0.564 | 0.007 | 0.023 | 0.674 | 0.009 | 0.024 |
| | | | | (0.027) | (0.025) | | (0.028) | (0.027) |
| | | 50% | 0.592 | -0.001 | 0.017 | 0.703 | 0.006 | 0.022 |
| | | | | (0.028) | (0.027) | | (0.030) | (0.029) |
| H15 | 0.75 | 10% | 0.548 | 0.012 | 0.028 | 0.656 | 0.013 | 0.036 |
| | | | | (0.025) | (0.025) | | (0.027) | (0.025) |
| | | 30% | 0.564 | 0.012 | 0.023 | 0.674 | 0.013 | 0.038 |
| | | | | (0.027) | (0.026) | | (0.028) | (0.025) |
| | | 50% | 0.592 | 0.010 | 0.017 | 0.703 | 0.013 | 0.041 |
| | | | | (0.028) | (0.028) | | (0.030) | (0.027) |
| H16 | 1 | 10% | 0.547 | 0.014 | 0.053 | 0.656 | 0.014 | 0.049 |
| | | | | (0.025) | (0.023) | | (0.027) | (0.024) |
| | | 30% | 0.564 | 0.011 | 0.054 | 0.674 | 0.013 | 0.050 |
| | | | | (0.026) | (0.024) | | (0.028) | (0.024) |
| | | 50% | 0.592 | 0.007 | 0.056 | 0.703 | 0.010 | 0.052 |
| | | | | (0.028) | (0.025) | | (0.030) | (0.026) |

Simulation setting (H14)-(H16): sample size $n = 2000$, number of true mediators $m = 5$, number of potential mediators $p = 1000$, coefficients $a = 0.38$, $b = 0.5$, $r = 2.5$, and $l = 0.1$ for all true mediators. $g = (0.5, 0.75, 1)$ for simulation settings H14-16, separately. Abbreviations: g, coefficient of covariate Z; Censoring, censoring rate; $R^2_{mediated}$, $R^2$-like measure for mediation effect estimation using w method; partial$\widehat{R^2}_{mediated}$, bias of estimated partial $R^2_{mediated}$ by MASH, where the corresponding standard deviation presented in brackets; unadjust$\widehat{R^2}_{mediated}$, bias of estimated $R^2_{mediated}$ by MASH without adjusting confounding, where the standard deviation presented in brackets; $SOS$, $SOS$ measure for mediation effect estimation using w method; partial$\widehat{SOS}$, bias of estimated partial $SOS$ by MASH, where the standard deviation presented in brackets; unadjust$\widehat{SOS}$, bias of estimated $SOS$ by MASH without adjusting confounding, where the corresponding standard deviation presented in brackets.

Table S6: Results of all settings in variable selection

| censoring | setting | H1 | H2 | H3 | H4 | H5 | H6 | H7 | H8 |
|---|---|---|---|---|---|---|---|---|---|
| 10% | TP | 99.20% | 100.00% | 100.00% | 100.00% | 100.00% | 100.00% | 100.00% | 100.00% |
| | FP | 0.42% | 0.56% | 0.49% | 0.49% | 0.54% | 0.51% | 0.46% | 0.07% |
| 30% | TP | 99.60% | 100.00% | 100.00% | 100.00% | 100.00% | 100.00% | 100.00% | 100.00% |
| | FP | 0.42% | 0.52% | 0.50% | 0.49% | 0.52% | 0.52% | 0.45% | 0.07% |
| 50% | TP | 99.40% | 100.00% | 99.92% | 100.00% | 99.96% | 100.00% | 100.00% | 100.00% |
| | FP | 0.31% | 0.70% | 0.48% | 0.46% | 0.54% | 0.50% | 0.46% | 0.10% |
| censoring | setting | H9 | H10 | H11 | H12 | H13 | H14 | H15 | H16 |
| 10% | TP | 100.00% | 100.00% | 99.92% | 100.00% | 100.00% | 100.00% | 100.00% | 100.00% |
| | FP | 0.49% | 0.44% | 0.32% | 0.50% | 0.30% | 0.25% | 0.19% | 0.11% |
| 30% | TP | 100.00% | 99.98% | 98.62% | 100.00% | 100.00% | 100.00% | 100.00% | 100.00% |
| | FP | 0.50% | 0.40% | 0.13% | 0.60% | 0.30% | 0.24% | 0.19% | 0.12% |
| 50% | TP | 99.98% | 97.44% | 83.82% | 100.00% | 100.00% | 100.00% | 100.00% | 100.00% |
| | FP | 0.45% | 0.26% | 0.04% | 0.70% | 0.30% | 0.25% | 0.19% | 0.15% |

Variable selection performance of the 16 simulation settings in terms of TP and FP. Abbreviations: TP, true positive rate; FP, false positive rate.

Table S7: Results of simulation studies using data with a confounder under $R^2$-like measure

| n | Censoring | $R^2_{mediated}$ | SOS |
|---|---|---|---|
| 500 | 10% | 0.015 | 0.017 |
| | | (0.060) | (0.066) |
| | 30% | -0.035 | -0.040 |
| | | (0.128) | (0.146) |
| | 50% | -0.191 | -0.219 |
| | | (0.206) | (0.235) |
| 300 | 10% | -0.122 | -0.142 |
| | | (0.160) | (0.184) |
| | 30% | -0.245 | -0.282 |
| | | (0.192) | (0.220) |
| | 50% | -0.407 | -0.466 |
| | | (0.175) | (0.199) |

Small sample size simulation settings: sample size $n = 300$ and 500, number of true mediators $m = 10$, number of potential mediators $p = 1000$, coefficients $a = 0.275$, $b = 0.2$, and $r = 2.5$ for all true mediators. Abbreviations: $n$, sample size; Censoring, censoring rate; $R^2_{mediated}$, $R^2$-like measure for mediation effect estimation using w method; $SOS$, $SOS$ measure for mediation effect estimation using w method. For $R^2_{mediated}$ and $SOS$, the first row is the bias and the second row is the corresponding standard deviation in brackets.

### 4 LIN54

LIN54, a protein-coding gene, is a component of the LIN, or DREAM complex, an essential regulator of cell cycle genes [2]. Although this gene has not been reported to be associated with DLBCL in the literatures, we looked into the gene expression and copy number variation of LIN54 and found both of them are borderline significant for the overall survival of DLBCL patients.

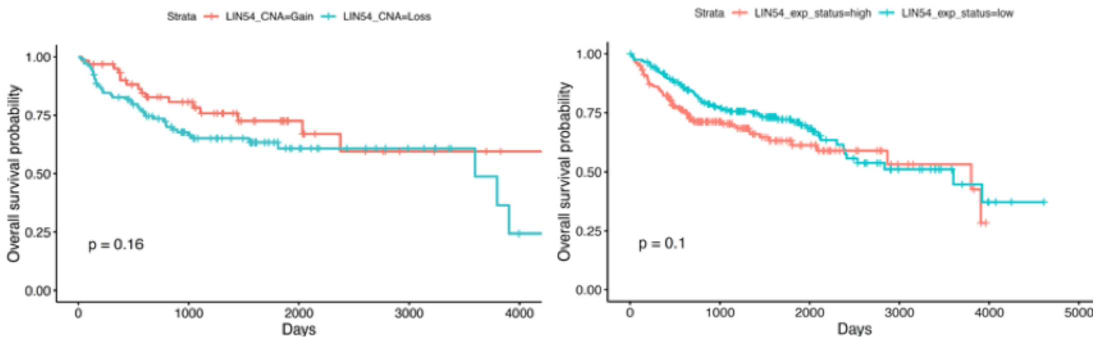


Figure 1: **Kaplan-Meier plot of overall survival of DLBCL for gene expression and copy number variation.** For expression data, we used median as cutoff to define the patients to LIN54 high and low groups. Similarly for CNV data, we set the medium value as cutoff for LIN54 gain and loss.

[1] KENT, J. T., and O'QUIGLEY, J. (1988). Measures of Dependence for Censored Survival Data. Biometrika 75(3) 525–4.

[2] Schmit, F., Cremer, S., Gaubatz, S. (2009). LIN54 is an essential core subunit of the DREAM/LINC complex that binds to the cdc2 promoter in a sequence-specific manner. The FEBS journal, 276(19), 5703–5716.
